# Chemoenzymatic Generation of Phospholipid Membranes Mediated by Type I Fatty Acid Synthase

**DOI:** 10.1101/2021.04.21.440816

**Authors:** Satyam Khanal, Roberto J. Brea, Michael D. Burkart, Neal K. Devaraj

## Abstract

The *de novo* formation of lipid membranes from minimal reactive precursors is a major goal in synthetic cell research. In nature, the synthesis of membrane phospholipids is orchestrated by numerous enzymes, including fatty acid synthases and membrane-bound acyltransferases. However, these enzymatic pathways are difficult to fully reproduce *in vitro*. As such, the reconstitution of phospholipid membrane synthesis from simple metabolic building blocks remains a challenge. Here, we describe a chemoenzymatic strategy for lipid membrane generation that utilizes a soluble bacterial fatty acid synthase (cgFAS I) to synthesize palmitoyl-CoA *in situ* from acetyl-CoA and malonyl-CoA. The fatty acid derivative spontaneously reacts with a cysteine-modified lysophospholipid by native chemical ligation (NCL), affording a non-canonical amidophospholipid that self-assembles into micron-sized membrane-bound vesicles. To our knowledge, this is the first example of reconstituting phospholipid membrane formation directly from acetyl-CoA and malonyl-CoA precursors. Our results demonstrate that combining the specificity and efficiency of a type I fatty acid synthase with a highly selective bioconjugation reaction provides a biomimetic route for the *de novo* formation of membrane-bound vesicles.

---

All living organisms use phospholipid membranes to control the exchange of materials with the extracellular matrix, isolate and protect sensitive chemical reactions, and maintain homeostasis inside cells.^1^ Furthermore, cells require phospholipid membranes for energy production, membrane protein synthesis, and cell signaling.^2,3^ Phospholipids are enzymatically generated by membrane-bound acyl-transferases as part of the Kennedy lipid synthesis pathway.^4,5^ The fatty acids needed for these biochemical reactions are in turn made by fatty acid synthases (FASs). FASs are ubiquitous proteins that catalyze the synthesis of fatty acids in a highly efficient manner through modularization of enzymatic functions and the use of carrier-mediated substrate shuttling.^6^ Through coordinated action, the different enzymatic modules of the FAS biosynthesize an elongated acyl chain of a specific length. Typical fatty acid products of FASs are palmitic, stearic and oleic acid.^7–9^

Drawing inspiration from the biochemical pathways for phospholipid synthesis, various research groups have devised strategies for the bottom-up construction of phospholipid membranes.^10–12^ Although recent studies have shown that phospholipids can be generated abiotically using chemical coupling reactions,^13,14^ the alkyl chains are typically synthesized in advance and provided externally. Currently, there are no known biomimetic strategies that synthesize phospholipids *de novo* by coupling *in situ* formed fatty acid tails with single-chain amphiphiles. Instead of externally adding alkyl species, we sought to harness a FAS for the formation of activated fatty acids from simple metabolic building blocks such as acetyl-CoA and malonyl-CoA. Subsequently, the activated fatty acids could chemically react with lipid precursors to form phospholipid membranes. Such a chemoenzymatic scheme would better mimic biological phospholipid synthesis compared to previous synthetic strategies and would enable the use of very short chain acyl-CoAs to drive membrane formation. Here, we employ a bacterial type I FAS (cgFAS I), in combination with native chemical ligation (NCL), to spontaneously generate membrane-forming synthetic phospholipids from simple water-soluble fatty acid precursors (Figure 1). Our chemoenzymatic approach enables *de novo* membrane formation, that is membrane formation in the absence of preexisting membranes. Chemoenzymatic phospholipid formation may provide simpler strategies to generate membrane compartments in synthetic cells,^10,13,15,16^ support the advancement of methods for reconstituting membrane proteins,^17,18^ and facilitate the synthesis of natural and non-canonical lipids.^19^

**Figure 1.**
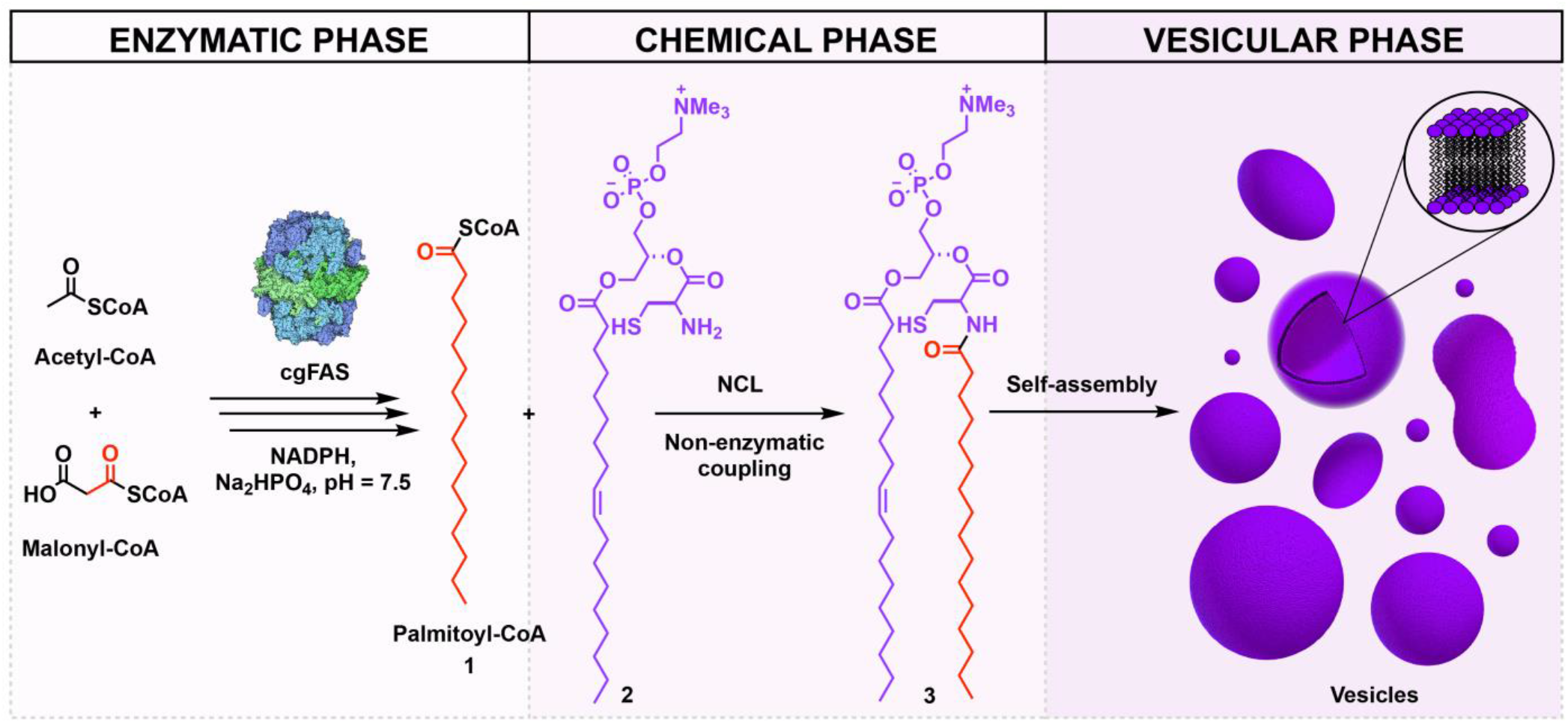
Schematic representation of FAS-mediated chemoenzymatic phospholipid synthesis. A bacterial type I FAS synthesizes palmitoyl-CoA **1** *in situ* using acetyl-CoA, malonyl-CoA and NADPH. **1** subsequently reacts with cysteine-modified lysophospholipid **2** via native chemical ligation (NCL) to form phospholipid **3**, which spontaneously self-assembles into membrane-bound vesicles.

We first identified an appropriate FAS for the *in situ* formation of activated fatty acids. While type II FASs are comprised of multiple enzymes working in a coordinated fashion, type I FASs consist of a single multi-enzyme complex with catalytic domains that interact with each other in a cooperative manner to form fatty acids.^6^ Through iterative cycles, a type I FAS utilizes acetyl-CoA and malonyl-CoA to produce medium-chain fatty acids in a stoichiometric fashion (Figure 2A). We considered that a type I FAS would be an ideal enzyme for our system as it would require reconstituting a single multi-domain protein. Additionally, some type I FASs, such as yeast and bacterial FASs, produce fatty acyl-CoA as their final product.^7^ As coenzyme A is a good leaving group, we reasoned that the bacterial type I FAS would be an appropriate enzyme for generating activated fatty acid products that could be efficiently coupled with appropriate thioester-reactive lysophospholipids to form non-canonical phospholipids *in situ*. We chose to work with type I FAS B from *Corynebacterium glutamicum* (cgFAS I) as it has been shown to primarily produce palmitoyl-CoA^20^ (Figure 2A) and has been efficiently expressed in *E. coli*.^21^

**Figure 2.**
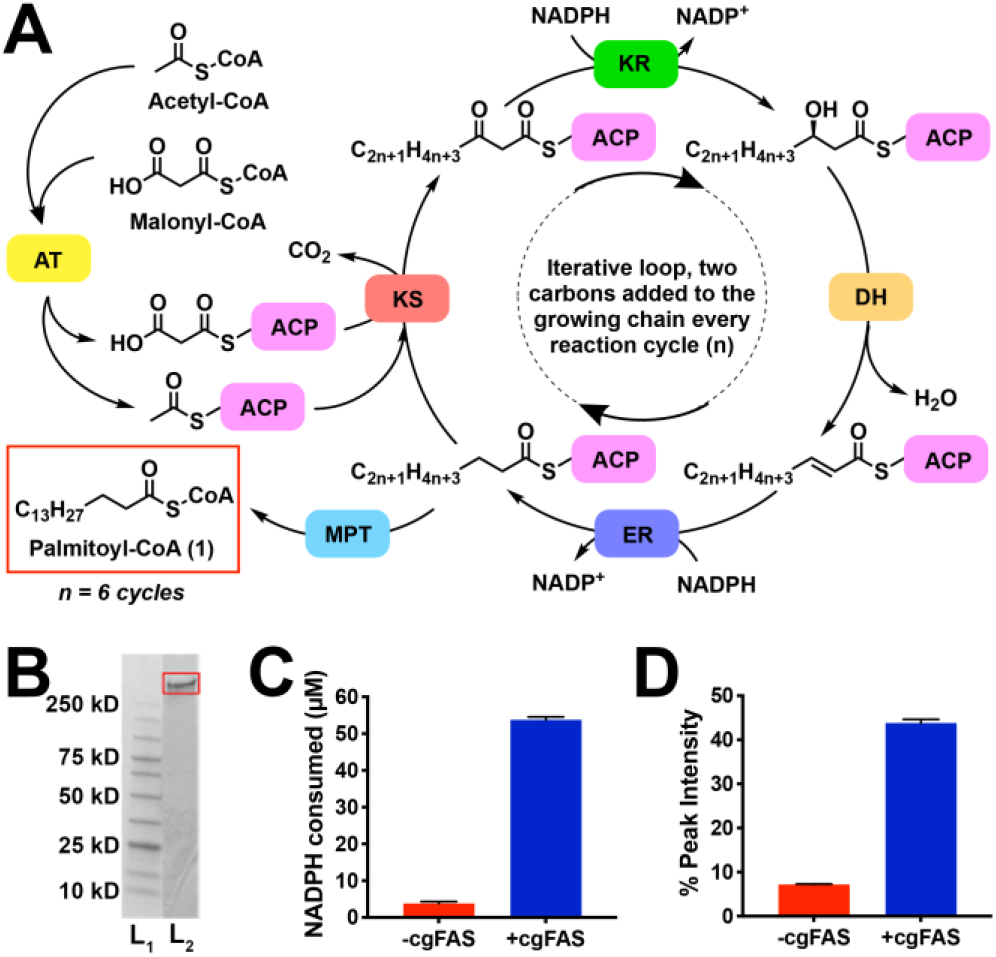
*In situ* synthesis of palmitoyl-CoA **1** mediated by bacterial type I FAS (cgFAS). A) Schematic representation of the iterative fatty acid elongation cycle. Malonyl-palmitoyltransferase (MPT) transfers the final palmitoyl moiety to a CoA molecule to form **1** [AT: acyl transferase; ACP: acyl carrier protein; KS: ketosynthase; KR: ketoreductase; DH: dehydratase; ER: enoyl reductase]. B) SDS-PAGE analysis of the His-tagged cgFAS after FPLC purification. Lane 1 (L_1_): ladder; Lane 2 (L_2_): purified His-tagged cgFAS (325 kDa). C) NADPH consumption assay analysis, verifying cgFAS activity. D) GC-MS FAME analysis of the cgFAS-catalyzed formation of **1** over 1 h.

N-terminal His_6_-tagged type I cgFAS was expressed in *E. coli* and purified by adapting a previously published procedure.^8^ The integrity of the protein, as well as its oligomeric state, were verified using size exclusion chromatography (SEC) on a fast protein liquid chromatography (FPLC) column (Figure 2B). The fractions were then subjected to a nicotinamide adenine di-nucleotide phosphate (NADPH) consumption assay to verify cgFAS activity (Figure 2C). First, we treated cgFAS I (100 nM) with acetyl-CoA (100 μM), malonyl-CoA (700 μM) and NADPH (1 mM) in 1 mM phosphate (Na_2_HPO_4_/NaH_2_PO_4_) buffer, pH 7.4 containing 1 mM tris(2-carboxyethyl)phosphine hydrochloride (TCEP) at 37 °C. Subsequently, we monitored NADPH oxidation to NADP^+^ over time by the decrease in fluorescence at 470 nm. The amount of NADPH consumed was verified by making a calibration curve using commercially available NADPH (Figure S2B). In agreement with previous reports, we observed that palmitoyl-CoA **1** was the major product of the cgFAS I-catalyzed reaction.^20,21^ Using gas chromatography-mass spectrometry (GC-MS) after fatty acid methyl ester (FAME) formation (Figure 2D, Figure S3), we observed that **1** comprised 93% of the total fatty acid species formed. Moreover, using the NADPH consumption assay together with high performance liquid chromatography-mass spectrometry (HPLC-MS), we observed that 41 μM of **1** was produced by the cgFAS I-mediated reaction, corresponding to 40.6% yield.

We next proceeded to select an appropriate thioester-reactive lysophospholipid for chemical coupling with cgFAS I-synthesized palmitoyl-CoA. We had previously prepared a novel class of cysteine-modified lysophospholipids that can undergo spontaneous acylation by NCL reaction with long-chain thioesters.^10,16,22^ We therefore hypothesized that cysteine-modified lysolipids would react by NCL with palmitoyl-CoA **1** generated *in situ* by cgFAS I. As a test, we synthesized cysteine-modified lysophospholipid **2** (Scheme S1A) and demonstrated NCL coupling with commercially available palmitoyl-CoA, forming phospholipid **3** (Figure S4B). Briefly, we treated lysophospholipid **2** (1 mM) with palmitoyl-CoA (1 mM) in 10 mM phosphate (Na_2_HPO_4_/NaH_2_PO_4_) buffer, pH 7.4 containing 10 mM TCEP at 37 °C. Phospholipid formation was followed using HPLC-MS combined with evaporative light-scattering detection (ELSD) and corroborated by chemically characterized standards (Scheme S1B, Figure S4). Using calibration curves, we determined that 820 μM of phospholipid **3** was formed after 3 h, corresponding to a yield of 82%.

In previous work, we have observed that amphiphilic species are preferentially acylated by amphiphilic reactants, likely promoted by co-assembly in micelles or membranes.^13,15,22^ To better understand the role self-assembly plays in the formation of the phospholipid product, we investigated the reactivity of lysophospholipid **2** with non-amphiphilic small-chain thioesters. Malonyl- and acetyl-CoA were selected as reactive thioester partners with **2**. Although both substrates contain a reactive thioester moiety that can react by NCL with cysteine-modified lysophospholipid **2**, the absence of a long-chain hydrophobic tail precludes assembly into structures such as micelles. Therefore, we anticipated a difference in their reactivity with **2** in comparison to the previously tested palmitoyl-CoA **1**. As expected, when we attempted to react **2** with malonyl- or acetyl-CoA under our standard NCL reaction conditions, we were unable to detect product formation (Figure S5).

To determine the ability of non-canonical phospholipid **3** to form membrane-bound vesicles, microscopy studies were performed. Neither palmitoyl-CoA **1** nor lysophospholipid **2** formed membranes in aqueous solution. On the other hand, phospholipid **3** readily formed membrane-bound assemblies when hydrated (Figure 3). Lipid vesicles were initially identified by phase-contrast (Figure 3A) and fluorescence microscopy using the membrane-staining dye BODIPY-FL DHPE (Figure 3B, Figure S6A). Under these conditions, vesicles of 1-10 μm diameter were observed after hydration and tumbling of **3** in phosphate buffer, pH 7.4 at 37 °C for 1 h. Transmission electron microscopy (TEM) also corroborated the formation of vesicular structures (Figure 3C). The encapsulation ability of the phospholipid vesicles was demonstrated by hydrating a thin lipid film of **3** in the presence of 8-hydroxypyrene-1,3,6-trisufonic acid (HPTS), a highly polar fluorescent dye, followed by removal of excess dye by spin-filtration and vesicle characterization using fluorescence microscopy (Figure 3D, Figure S6B).

**Figure 3.**
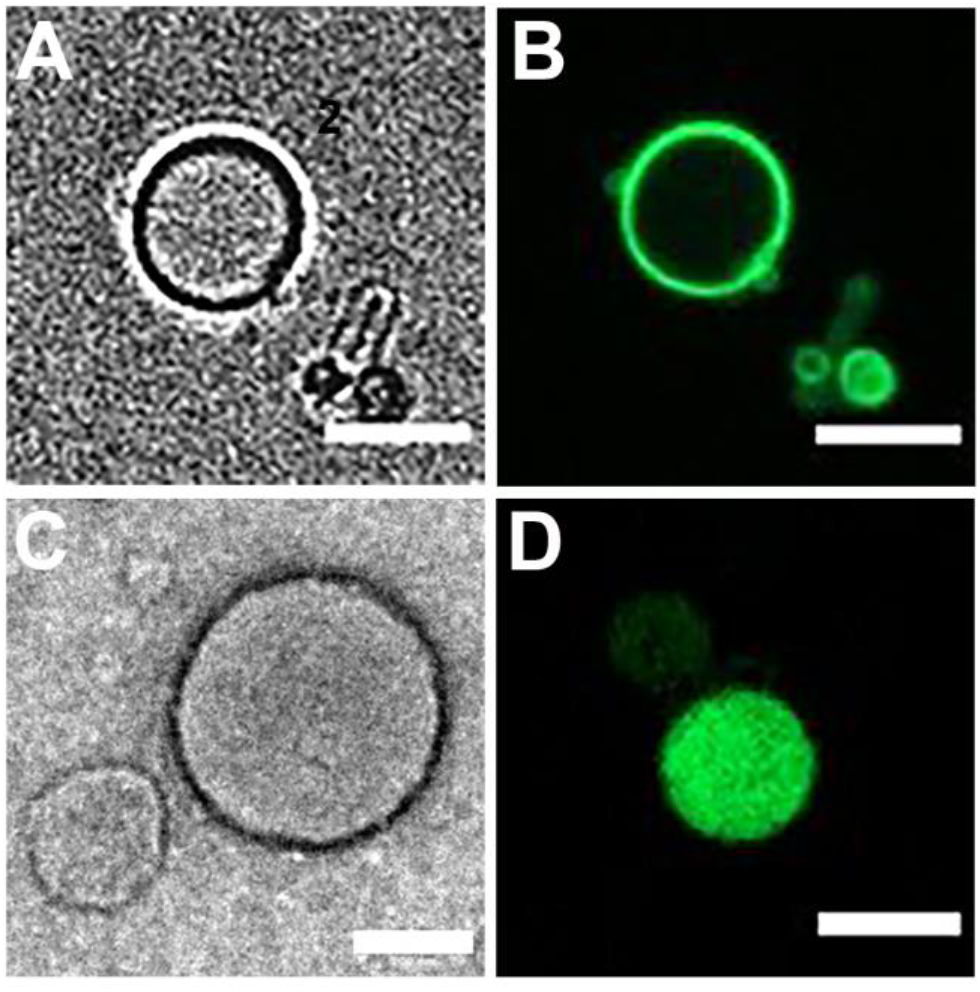
Characterization of phospholipid **3** vesicular structures. A) Phase-contrast microscopy image of membrane-bound vesicles resulting from the self-assembly of **3**. Scale bar denotes 5 μm. B) Fluorescence microscopy image of vesicles formed by hydration of a thin film of **3.** Membranes were stained with 0.1 mol % BODIPY-FL DHPE. Scale bar denotes 5 μm. C) TEM image of negatively stained vesicles of **3**. Scale bar denotes 100 nm. D) Fluorescence microscope image demonstrating the encapsulation of HPTS in vesicles of **3**. Scale bar denotes 5 μm.

Having characterized the individual enzymatic and chemical reactions, we next explored combining enzymatic palmitoyl-CoA **1** synthesis with chemical phospholipid **3** synthesis in a one-pot reaction (Figure 4). Briefly, we added lysophospholipid **2** (400 μM) to 10 mM phosphate (Na_2_HPO_4_/NaH_2_PO_4_) buffer, pH 7.4 containing cgFAS I (1 μM), acetyl-CoA (1 mM), malonyl-CoA (1 mM) and NADPH (10 mM) along with 10 mM (TCEP) at 37 °C. Phospholipid formation was followed using HPLC-MS-ELSD measurements. Optimization of the reaction conditions enabled rapid coupling between cgFAS I generated **1** and **2**. The one-pot reaction afforded the corresponding phospholipid **3** as the prominent product within 30 min (Figure 4A). All of lysophospholipid **2** was consumed in less than 4 h (Figure 4 B) to afford 367 μM of phospholipid **3**. After 30 min of reaction, small vesicular structures were detected by fluorescence microscopy using BODIPY-FL DHPE (Figure S7A). After leaving the reaction tumbling overnight at 37 °C, we observed larger vesicles in the range of 1-2 μm in diameter (Figure 4C, Figure S7B).

**Figure 4.**
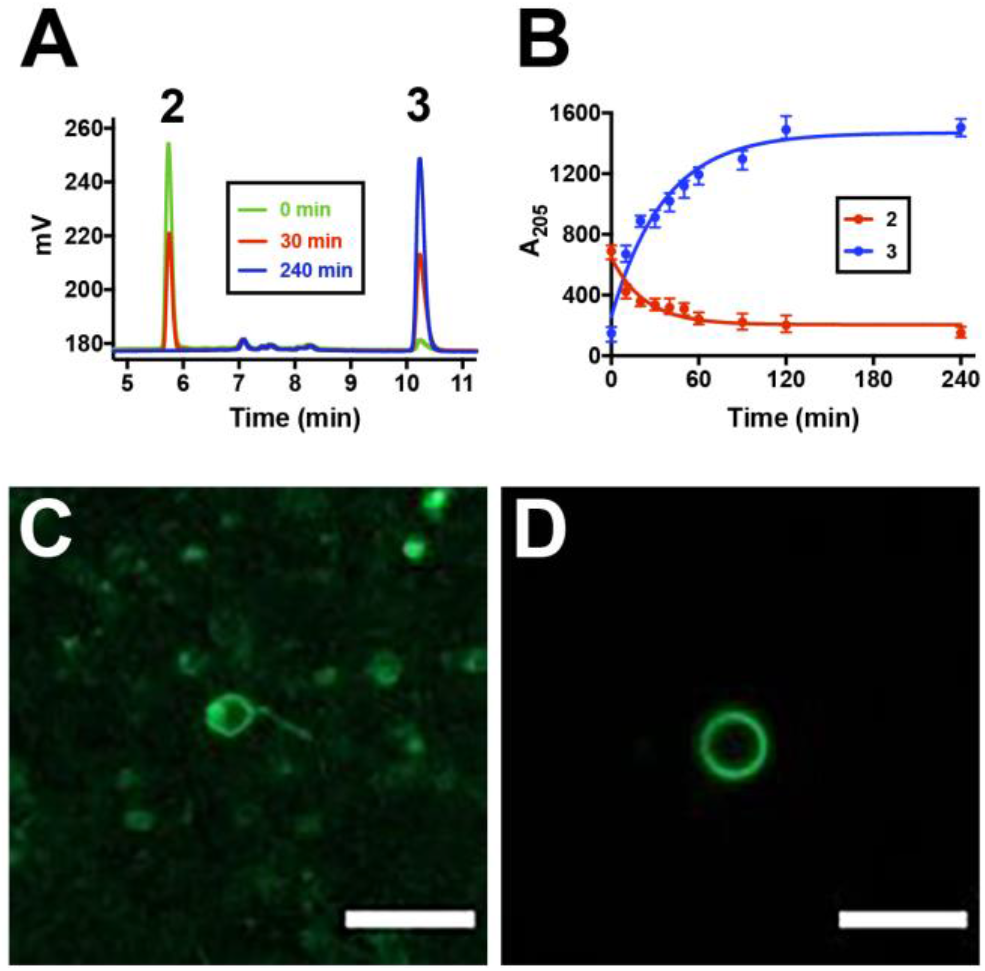
One-pot chemoenzymatic formation and self-assembly of phospholipid **3**. A) HPLC traces corresponding to the NCL-based synthesis of phospholipid **3** from enzymatically generated **1** and lysophospholipid **2.** B) Kinetic curves of lysophospholipid **2** consumption and phospholipid **3** formation during one-pot synthesis. C) Fluorescence microscopy image of phospholipid **3** vesicles after 4 h of chemoenzymatic reaction. D) Fluorescence microscopy image of chemoenzymatically formed vesicles of **3** in the presence of GuHCl, decanol and cholesterol. Membranes were stained with 0.1 mol % BODIPY-FL DHPE. Scale bar denotes 5 μm.

We next investigated the one-pot chemoenzymatic formation of membranes in the presence of biologically relevant cell membrane components, including cholesterol^23^, ionic small molecules such as guanidine hydrochloride (GuHCl)^24–26^, and short-chain alkanols such as decanol^27^. We added cholesterol (400 μM), GuHCl (400 μM) and 1-decanol (400 μM) to the one-pot *in situ* chemoenzymatic reaction forming phospholipid **3**. We observed that the additives did not perturb the formation of phospholipid **3** membranes, and, if anything, led to the formation of larger vesicles. Vesicles were stable over 48 h at 37 °C, as observed by fluorescence microscopy using BODIPY-FL DHPE (Figure 4D).

In summary, we have developed a chemoenzymatic route to synthesize non-canonical phospholipids from water-soluble precursors. We showed that palmitoyl-CoA generated *in situ* by bacterial type I FAS effectively couples with a cysteine-modified lysophospholipid. The resulting phospholipid synthesis triggers *de novo* formation of membrane-bound vesicles. Given our approach, there should be flexibility to diversify the lipid species generated in the reaction. Even though we utilized cgFAS I to selectively produce palmitoyl-CoA, the use of fatty acid synthases from other organisms could enable the formation of a diverse array of fatty acyl-CoA species, which could be subsequently coupled to reactive lysophospholipids to give several non-canonical lipid species. For instance, many bacterial FAS are known to synthesize terminally branched iso-, anteiso-, or omega-alicyclic fatty acids from branched, short-chain carboxylic acid precursors such as methylmalonyl-CoA.^7,28,29^ We plan on utilizing the *in situ* synthesis of diverse phospholipid species to facilitate investigations of how lipid membrane composition affects vesicle assembly, growth and division.

## AUTHOR INFORMATION

### Notes

The authors declare no competing financial interest.

## ACKNOWLEDGMENT

This material is based upon work supported by the National Science Foundation (EF-1935372) and the Department of Defense (Army Research Office) through the MURI program (W911NF-13-1-0383). We sincerely thank Prof. Jay Keasling (University of California, Berkeley) for generously providing us the plasmid for the type I FAS B from *Corynebacterium glutamicum* (cgFAS I). We thank Prof. Itay Budin (University of California, San Diego) and Prof. Martin Grininger (Goethe University Frankfurt) for helpful discussions. Satyam Khanal acknowledges support from a Roger Tsien Fellowship through UCSD.

