## Supplementary Information for "Chemoenzymatic Generation of Phospholipid Membranes Mediated by Type I Fatty Acid Synthase"

**Table of Contents**

1. **General Methods, Instrument Details and Materials**
2. **Experimental Procedures**
3. **Supplementary Schemes**
4. **Supplementary Figures**
5. **NMR Spectra**
6. **References**

### 1. General Methods, Instrument Details and Materials

Commercially available 1-oleoyl-2-hydroxy-*sn*-glycero-3-phosphocholine (Lyso $C_{18:1}$ PC-OH) was used as obtained from Avanti® Polar Lipids. *N*-Boc-*L*-Cys(Trt)-OH, *N*-Fmoc-*L*-Cys(Trt)-OH, 2,4,6-trichlorobenzoyl chloride (TCBC), 4-dimethylaminopyridine (DMAP), *N,N'*-diisopropylcarbodiimide (DIC), O-(7-azabenzotriazol-1-yl)-1,1,3,3-tetramethyl-uronium hexafluorophosphate (HATU), dichloromethane ($CH_2Cl_2$), *N*,*N*-dimethylformamide (DMF), *N*,*N*-diisopropylethylamine (DIEA), trifluoroacetic acid (TFA), triethylsilane (TES), 4-methylpiperidine, palmitic acid and tris(2-carboxyethyl)phosphine hydrochloride (TCEP) were obtained from Sigma-Aldrich. Texas Red® 1,2-dihexadecanoyl-*sn*-glycero-3-phosphoethanolamine, triethylammonium salt (Texas Red® DHPE) was obtained from Life Technologies. BODIPY-FL DHPE was obtained from ThermoFisher Scientific. Deuterated chloroform ($CDCl_3$) and methanol ($CD_3OD$) were obtained from Cambridge Isotope Laboratories. All reagents obtained from commercial suppliers were used without further purification unless otherwise noted. Solvent mixtures for chromatography are reported as v/v ratios. HPLC analysis was carried out on an Eclipse Plus C8 analytical column with *Phase A*/*Phase B* gradients [*Phase A*: $H_2O$ with 0.1% formic acid; *Phase B*: MeOH with 0.1% formic acid]. HPLC purification was carried out on Zorbax SB-C18 semipreparative column with *Phase A*/*Phase B* gradients [*Phase A*: $H_2O$ with 0.1% formic acid; *Phase B*: MeOH with 0.1% formic acid]. GC-MS analysis was carried out on an Agilent 7890A GC system connected to a 5975C VL MSD quadrupole MS (EI). Samples were separated on a 60m DB23 Agilent GCMS column using helium as carrier gas and a gradient of 110 °C to 200 °C at 15 °C/min, followed by 20 min at 200 °C. Proton nuclear magnetic resonance ($^1H$ NMR) spectra were recorded on a Varian VX-500 MHz spectrometer, and were referenced relative to residual proton resonances in $CDCl_3$ (at δ 7.24 ppm) or $CD_3OD$ (at δ 4.87 or 3.31 ppm). Chemical shifts were reported in parts per million (ppm, δ) relative to tetramethylsilane (δ 0.00). $^1H$ NMR splitting patterns are assigned as singlet (s), doublet (d), triplet (t), quartet (q) or pentuplet (p). All first-order splitting patterns were designated on the basis of the appearance of the multiplet. Splitting patterns that could not be readily interpreted are designated as multiplet (m) or broad (br). Carbon nuclear magnetic resonance ($^{13}C$ NMR) spectra were recorded on a Varian VX-500 MHz spectrometer, and were referenced relative to residual proton resonances in $CDCl_3$ (at δ 77.23 ppm) or $CD_3OD$ (at δ 49.15 ppm). Electrospray Ionization-Time of Flight (ESI-TOF) spectra were obtained on an Agilent 6230 Accurate-Mass TOF-MS mass spectrometer. Spinning-disk confocal microscopy images were acquired on a Yokagawa spinning-disk system (Yokagawa, Japan) built around an Axio Observer Z1motorized inverted microscope (Carl Zeiss Microscopy GmbH, Germany) with a 63x, 1.40 NA oil immersion objective to an Evolve 512x512 EMCCD camera (Photometrics, Canada) using ZEN imaging software (Carl Zeiss Microscopy GmbH, Germany). A condenser/objective with a phase stop of Ph2 was used to obtain the phase-contrast images. The fluorophores were excited with a 20 mW DPSS laser (Texas Red®). NanoDrop 2000C spectrophotometer was used for UV/Vis measurements. Fluorescence measurements were carried out on a Tecan infinite F200 plate reader instrument. Transmission electron microscopy (TEM) images were recorded on a FEI Tecnai™ Sphera 200 kV microscope equipped with a $LaB_6$ electron gun, using the standard cryotransfer holders developed by Gatan, Inc.

### 2. Experimental Procedures

Expression and purification of cgFAS

pBbE5c (a pET22b[+] derivative) was kindly provided by Dr. Robert Haushalter from Professor Dr. Jay Keasling's lab in University of Berkeley.[1] This plasmid contained the *Corynebacterium* type I FAS gene [codon optimized]. It has been shown that *Corynebacteria* have two type I FASs, type IA and type I B[2]. Type I A is the essential FAS since it makes the bulk of the fatty acid whereas type I B variant supplements palmitoyl-CoA. For this project, we required *in situ* palmitoyl-CoA formation, so we utilized type I B of *Corynebacterium*. We then added a C-terminal histidine $His_6$-tag onto the plasmid using forward and reverse primers for $His_6$-tag. The ACP domain of type I bacterial FASs requires activation by an external phosphopantetheine transferase (sfp) that adds a phosphopantetheine arm onto a conserved serine residue on the ACP. Dr. Haushalter also provided us with *Escherichia coli* having the phosphopantetheine transferase (Sfp) embedded into its genome and we made competent cells out of the provided sample in order to express the cgFAS gene with an activated ACP domain. For making competent *E. coli* cells, we used an established protocol provided by New England Biolabs (NEB).[3] The pBbE5c plasmid was then transformed into the competent *E. coli* cells and grown overnight at 37 °C in Luria-Bertani (LB) broth containing 0.1 mg/mL of kanamycin. Afterwards, 1 mL of the overnight culture was used to inoculate 1 L of freshly autoclaved LB medium containing 0.1 mg/mL of kanamycin. The rest of the overnight culture was stored as 25% glycerol stocks at -80 °C. The culture was grown at 37 °C in a shaker-incubator till the $OD_{600}$ reached 0.6. Overexpression of cgFAS was induced by addition of 1 mM isopropyl 1-thio-D-galactopyranoside (IPTG). The cells were then grown for 16 h at 18 °C, after which they were harvested by centrifuging at 6,000 rcf for 20 min at 4 °C. The pellet was resuspended by vortexing in 10 mL of lysis buffer containing 25 mM Tris buffer, pH 8.0, 0.5 M NaCl, 2 mM β-mercaptoethanol, 1 mg/mL lysozyme and a cocktail of protease inhibitors (SigmaFast®). Following cell lysis by an ultrasonicator probe, debris were removed by centrifuging (13,000 rcf, 20 min, 4 °C). The supernatant was incubated in a gravity column with $Ni^{2+}$-nitrilotriacetate (Ni-NTA) agarose resin pre-equilibrated with 10 mM imidazole on a shaker table for 3 h at 4 °C. Next, the flow-through was discarded and the resin was washed with 10 mL each of 20 mM imidazole and 50 mM imidazole solutions. Finally, $His_6$-tagged cgFAS was eluted with 250 mM imidazole and collected in 1 mL fractions. The fractions were analyzed by SDS-PAGE to check for considerable impurities, and subsequently concentrated in a centrifugal filter with a 100,000 nominal molecular weight limit (Amicon Ultra-4, Merck Millipore). The sample was further purified by size-exclusion chromatography (Fig. 2B) (column: Superose 6 10/300GL, GE Healthcare, buffer G: 100 mM $Na_2HPO_4/NaH_2PO_4$ pH 7.4, 100 mM NaCl) and examined for its oligomeric state. Fractions were pooled, concentrated and aliquoted to a final concentration of 1 µM of protein containing 10% glycerol each and stored at -80 °C. Protein $A_{280}$ and concentration measurements carried out using a NanoDrop (ThermoFisher Scientific).

cgFAS I activity assay

To test the activity of cgFAS I, NADPH consumption assay was carried out on a 50 µL scale, including 35 µL of 100 mM $NaH_2PO_4/Na_2HPO_4$ buffer, pH 7.4 (containing 10 mM TCEP), 5 µL of 1 mM acetyl-CoA (in sterile $H_2O$), 5 µL of 10 mM NADPH (in sterile $H_2O$) and 5 µL of cgFAS I (1 µM in elution buffer) at 37 °C. After 1 min of recording the emission at 470 nm using a Spark multimode plate reader (Tecan), the reaction was started by the addition of 5 µL of 7 mM malonyl-CoA and the emission was continuously measured for 1 h (Fig. S2A). NADPH consumption was closely followed, and a calibration curve was made to observe the amount of NADPH consumed in the reaction (Fig. S2B). After subtracting non-specific decay of signal, we observed that 602 µM of NADPH was being consumed in the reaction, which correlates to 43 µM fatty acid species being made in the reaction.

GC-MS FAME analysis of FAS product

The product from the cgFAS I activity assay was subjected to GC-MS fatty acid methyl ester (FAME) analysis to investigate the fatty acid product. After 1 h of NADPH consumption assay, the reaction was centrifuged to remove protein debris and the sample was then resuspended in 300 µL of 1 M methanolic acid and incubated at 65 °C for 30 min. The FAMEs were extracted using 300 µL of hexanes and then separated by GC-MS. As expected, the major peak was the methyl palmitate (Fig. 2D, Fig. S3). Using area under the curve, we observed that 93% of the fatty acid species was palmitoyl-CoA (**1**), with a small peak (4%) of stearoyl-CoA. The peak was corroborated using commercially available palmitoyl-CoA (100 nM final concentration) after conversion into FAME. We normalized the FAS-mediated reaction product peak to that of the commercially available methyl palmitate. We also ran a negative control alongside the reaction to ascertain the baseline spectrum in the absence of cgFAS, and accounted for this while calculating the final concentration of the FAS-mediated product (Fig. S2C).

HPLC-MS quantification of palmitoyl-CoA (**1**) product

Chromatographic separation was performed with a mobile-phase system with gradients based on *Phase C* (water, 10 mM triethylamine/acidic acid buffer, adjusted to pH 9.0) and *Phase D* (acetonitrile). A multistep gradient at a flow rate of 0.25 ml/min was used with the starting condition of *Phase D* at 7%, a linear increase to 60% until 6 min, then to 70% until 9.5 min and finally to 90% until 10 min runtime. cgFAS I-mediated palmitoyl-CoA (**1**) formation was set up as described in the activity assay and monitored over 1 h using HPLC-MS (Fig. S2C). For each HPLC-MS run, a small aliquot (10 µL) of the reaction was taken and centrifuged to remove protein debris before loading onto the column. For the quantification, the UV trace at 205 nm was used (Fig S2C). The amount of palmitoyl-CoA made was 41 µM, which closely agreed with the amount previously calculated from NADPH consumption (43 µM).

Synthesis of Lysophospholipids

Lysophospholipid **2** was synthesized according the Scheme S1A.

**1-oleoyl-2-[*N*-Boc-*L*-Cys(Trt)]-*sn*-glycero-3-phosphocholine (4).**[4] A solution of 1-oleoyl-2-hydroxy-*sn*-glycero-3-phosphocholine (**Lyso $C_{18:1}$ PC-OH**, 25.0 mg, 47.9 µmol), *N*-Boc-*L*-Cys(Trt)-OH (55.5 mg, 119.8 µmol), DMAP (35.1 mg, 287.5 µmol) and $Et_3N$ (23.4 µL, 167.7 µmol) in $CDCl_3$ (1.875 mL) was stirred at r.t. for 10 min. Then, TCBC (48.7 µL, 311.5 µmol) was added. After 12 h stirring at r.t., $H_2O$ (125 µL) was added to quench the acid chloride, and the solvent was removed under reduced pressure to give a pale yellow solid. The corresponding residue was dissolved in MeOH (1 mL), filtered using a 0.2 µm syringe-driven filter, and the crude solution was purified by HPLC, affording 39.6 mg of lysophospholipid **4** as a white solid [86%, $t_R$ = 7.8 min (Zorbax SB-C18 semipreparative column, 100% *Phase B*, 15.5 min)]. $^1$H NMR ($CDCl_3$, 500.13 MHz, δ): 7.38 (d, *J* = 7.5 Hz, 6H, 6 × $CH_{Ar}$), 7.33-7.26 (m, 6H, 6 × $CH_{Ar}$), 7.25-7.19 (m, 3H, 3 × $CH_{Ar}$), 5.40-5.29 (m, 2H, 2 × CH), 5.27-5.13 (m, 1H, 1 × CH), 5.07 (d, *J* = 9.0 Hz, 1H, 1 × NH), 4.41-3.88 (m, 7H, 3 × $CH_2$ + 1 × CH), 3.77-3.57 (m, 2H, 1 × $CH_2$), 3.25 (s, 9H, 3 × $CH_3$), 2.74-2.48 (m, 2H, 1 × $CH_2$), 2.31-2.09 (m, 2H, 1 × $CH_2$), 2.07-1.93 (m, 4H, 2 × $CH_2$), 1.61-1.43 (m, 2H, 1 × $CH_2$), 1.42 (s, 9H, 3 × $CH_3$), 1.31-1.17 (m, 20H, 10 × $CH_2$), 0.88 (t, *J* = 7.0 Hz, 3H, 1 × $CH_3$). $^{13}$C NMR ($CDCl_3$, 125.77 MHz, δ): 173.5, 170.4, 163.9, 155.3, 144.4, 130.1, 129.9, 129.7, 129.6,

128.2, 128.2, 127.1, 80.1, 72.3, 67.2, 66.5, 63.8, 62.7, 59.4, 54.6, 52.7, 34.1, 34.1, 34.0, 32.0, 29.9, 29.9, 29.7, 29.5, 29.5, 29.4, 29.4, 29.3, 29.2, 29.2, 28.5, 28.5, 27.4, 27.3, 24.9, 24.8, 22.8, 14.3. MS (ESI-TOF) [m/z (%)]: 989 ($[M+Na]^+$, 20), 967 ($[MH]^+$, 100). HRMS (ESI-TOF) calculated for $C_{53}H_{80}N_2O_{10}PS$ ($[MH]^+$) 967.5266, found 967.5269.

*Alternative method:*[5] A solution of *N*-Boc-*L*-Cys(Trt)-OH (88.9 mg, 191.7 µmol) in $CH_2Cl_2$ (7.5 mL) was stirred at rt for 10 min, and then DIC (45.0 µL, 287.5 µmol) and DMAP (11.7 mg, 95.8 µmol) were successively added. After 10 min stirring at rt, 1-oleoyl-2-hydroxy-*sn*-glycero-3-phosphocholine (25.0 mg, 47.9 µmol) was added. After 12 h stirring at rt, the solvent was removed under reduced pressure, and the crude was purified by HPLC, affording 34.6 mg of **3** as a colorless foam [75%].

**1-oleoyl-2-(*L*-Cys)-*sn*-glycero-3-phosphocholine (2).** A solution of 1-oleoyl-2-[N-Boc-*L*-Cys(Trt)]-*sn*-glycero-3-phosphocholine (**3**, 5.0 mg, 5.2 µmol) in 200 µL of TFA/$CH_2Cl_2$/TES (90:90:20) was stirred at rt for 30 min. After removal of the solvent, the residue was dried under high vacuum for 3 h. Then, the corresponding residue was dissolved in MeOH (500 µL), filtered using a 0.2 µm syringe-driven filter, and the crude solution was purified by HPLC, affording 3.1 mg of the lysophospholipid **2** as a colorless foam [82%, $t_R$ = 8.6 min (Zorbax SB-C18 semipreparative column, 50% *Phase A* in *Phase B*, 5 min, and then 5% *Phase A* in *Phase B*, 10 min)]. $^1H$ NMR ($CD_3OD$, 500.13 MHz, δ): 5.46-5.26 (m, 3H, 3 × CH), 4.49-4.35 (m, 1H, 1 × CH), 4.34-4.19 (m, 3H, 1.5 × $CH_2$), 4.16-3.99 (m, 3H, 1.5× $CH_2$), 3.72-3.59 (m, 2H, 1 × $CH_2$), 3.29-3.26 (m, 1H, 0.5× $CH_2$), 3.23 (s, 9H, 3 × $CH_3$), 3.19-3.00 (m, 1H, 0.5× $CH_2$), 2.35 (t, *J* = 6.7 Hz, 2H, 1 × $CH_2$), 2.09-1.90 (m, 4H, 2 × $CH_2$), 1.69-1.53 (m, 2H, 1 × $CH_2$), 1.41-1.22 (m, 20H, 10 × $CH_2$), 0.90 (t, *J* = 6.7 Hz, 3H, 1 × $CH_3$). $^{13}C$ NMR ($CD_3OD$, 125.77 MHz, δ): 174.9, 172.3, 131.6, 130.8, 73.7, 67.4, 64.9, 63.4, 63.2, 60.6, 54.6, 34.9, 33.7, 33.1, 30.8, 30.8, 30.7, 30.6, 30.5, 30.4, 30.3, 30.3, 30.2, 28.1, 26.0, 23.8, 14.5. MS (ESI-TOF) [m/z (%)]: 625 ($[MH]^+$, 100). HRMS (ESI-TOF) calculated for $C_{29}H_{58}N_2O_8PS$ ($[MH]^+$) 625.3651, found 625.3647.

Synthesis of Phospholipids

Phospholipid **3** was synthesized according to the Scheme S1B. The pure compound was used to corroborate by HPLC/ELSD/MS the formation of **3** using our NCL-FAS I approach.

**1-oleoyl-2-[*N*-Fmoc-*L*-Cys(Trt)]-*sn*-glycero-3-phosphocholine (5).** A solution of 1-oleoyl-2-hydroxy-*sn*-glycero-3-phosphocholine (**Lyso $C_{18:1}$ PC-OH**, 25.0 mg, 47.9 µmol), *N*-Fmoc-*L*-Cys(Trt)-OH (70.2 mg, 119.8 µmol), DMAP (35.1 mg, 287.5 µmol) and $Et_3N$ (23.4 µL, 167.7 µmol) in $CDCl_3$ (1.9 mL) was stirred at r.t. for 10 min. Then, TCBC (48.7 µL, 311.5 µmol) was added. After 12 h stirring at r.t., $H_2O$ (125 µL) was added to quench the acid chloride, and the solvent was removed under reduced pressure to give a pale yellow solid. The corresponding residue was dissolved in MeOH (500 µL), filtered using a 0.2 µm syringe-driven filter, and the crude solution was purified by HPLC, affording 42.7 mg of lysophospholipid **5** as a white solid [82%, $t_R$ = 7.8 min (Zorbax SB-C18 semipreparative column, 100% *Phase B*, 25 min)]. MS (ESI-TOF) [m/z (%)]: 1111 ($[M+Na]^+$, 100), 1089 ($[MH]^+$, 72).

**1-oleoyl-2-[*L*-Cys(Trt)]-*sn*-glycero-3-phosphocholine (6).** A solution of 1-oleoyl-2-[*N*-Fmoc-*L*-Cys(Trt)]-*sn*-glycero-3-phosphocholine (**5**, 7.5 mg, 6.9 µmol) in 125 µL of 4-methylpiperidine/$CH_2Cl_2$ (25:100) was stirred at rt for 30 min. After removal of the solvent, the residue was dried under high vacuum for 3 h. Then, the corresponding residue was dissolved in MeOH (250 µL), filtered using a 0.2 µm syringe-driven filter, and the crude solution was purified by HPLC, affording 5.2 mg of the lysophospholipid **6** as a colorless film [87%, $t_R$ = 8.3 min (Zorbax SB-C18 semipreparative column, 50-0% *Phase A* in *Phase B*, 2.5 min, and then 100% *Phase B*, 15 min)]. MS (ESI-TOF) [m/z (%)]: 889 ($[M+Na]^+$, 60), 867 ($[MH]^+$, 100).

**1-oleoyl-2-[*L*-Cys(Trt)-(palmitoyl)]-*sn*-glycero-3-phosphocholine (7).** A solution of palmitic acid (0.6 mg, 2.3 µmol) in $CH_2Cl_2$/DMF (1:1) (200 µL) was stirred at 0 ºC for 10 min, and then HATU (1.0 mg, 2.5 µmol) and DIEA (1.6 µL, 9.2 µmol) were successively added. After 10 min stirring at 0 ºC, 1-oleoyl-

2-[*L*-Cys(Trt)]-*sn*-glycero-3-phosphocholine (**6**, 2.0 mg, 2.3 µmol) was added. After 1 h stirring at rt, the mixture was concentrated under reduced pressure. The corresponding residue was dissolved in MeOH (250 µL), filtered using a 0.2 µm syringe-driven filter, and the crude solution was purified by HPLC, affording 2.0 mg of **7** as a colorless film [79%, $t_R$ = 25.0 min (Zorbax SB-C18 semipreparative column, 50-0% *Phase A* in *Phase B*, 2.5 min, and then 100% *Phase B*, 20 min)]. MS (ESI-TOF) [m/z (%)]: 1127 ($[M+Na]^+$, 37), 1105 ($[MH]^+$, 100).

**1-oleoyl-2-[*L*-Cys-(palmitoyl)]-*sn*-glycero-3-phosphocholine (3).** A solution of 1-oleoyl-2-[*L*-Cys(Trt)-(palmitoyl)]-*sn*-glycero-3-phosphocholine (**7**, 1.0 mg, 0.9 µmol) in 200 µL of TFA/$CH_2Cl_2$/TES (90:90:20) was stirred at rt for 30 min. After removal of the solvent, the residue was dried under high vacuum for 3 h. Then, the corresponding residue was dissolved in MeOH (250 µL), filtered using a 0.2 µm syringe-driven filter, and the crude solution was purified by HPLC, affording 0.7 mg of the amidophospholipid **3** as a colorless film [83%, $t_R$ = 16.4 min (Zorbax SB-C18 semipreparative column, 50-0% *Phase A* in *Phase B*, 2.5 min, and then 100% *Phase B*, 15 min)]. $^1$H NMR ($CD_3OD$, 500.13 MHz, δ): 8.54 (s, 1H, 1 × NH), 5.41-5.33 (m, 2H, 2 × CH), 5.32-5.22 (m, 1H, 1 × CH), 4.67-4.62 (m, 1H, 1 × CH), 4.46-4.36 (m, 1H, 0.5 × $CH_2$), 4.33-4.19 (m, 3H, 1.5 × $CH_2$), 4.12-3.96 (m, 2H, 1 × $CH_2$), 3.74-3.58 (m, 2H, 1 × $CH_2$), 3.23 (s, 9H, 3 × $CH_3$), 3.04-2.85 (m, 2H, 1 × $CH_2$), 2.34 (t, *J* = 7.5 Hz, 2H, 1 × $CH_2$), 2.30-2.23 (m, 2H, 1 × $CH_2$), 2.10-1.95 (m, 4H, 2 × $CH_2$), 1.71-1.53 (m, 4H, 2 × $CH_2$), 1.39-1.26 (m, 44H, 22 × $CH_2$), 0.90 (t, *J* = 6.8 Hz, 6H, 2 × $CH_3$). $^{13}$C NMR ($CD_3OD$, 125.77 MHz, δ): 176.4, 174.9, 171.0, 130.9, 130.8, 73.3, 67.4, 64.9, 63.5, 60.6, 56.2, 54.7, 36.7, 34.9, 33.1, 30.9, 30.9, 30.9, 30.8, 30.8, 30.8, 30.7, 30.7, 30.7, 30.6, 30.5, 30.5, 30.5, 30.4, 30.4, 30.4, 30.3, 30.3, 30.2, 28.2, 27.0, 26.7, 26.0, 23.8, 14.5. MS (ESI-TOF) [m/z (%)]: 863 ($[MH]^+$, 100). HRMS (ESI-TOF) calculated for $C_{45}H_{88}N_2O_9PS$ ($[MH]^+$) 863.5948, found 863.5947.

Rehydration of phospholipid **3**

5 µL of a 10 mM solution of purified phospholipid **3** in $CHCl_3$ was added to a 1 mL vial, placed under $N_2$ and dried for 15 min to prepare a lipid film. Then, 95 µL of $H_2O$ was added and the solution was tumbled at 25 ºC for 1 h. Afterwards, to 10 µL of this 500 µM aqueous solution of phospholipid **3** was added 0.1 µL of a 100 µM BODIPY-FL DHPE solution in EtOH, and the mixture was briefly agitated. The corresponding mixture was finally monitored by phase-contrast and fluorescence microscopy in order to determine the vesicle structure. We observed that the phospholipid product **3** spontaneously self-assemble into vesicles. Initially, the vesicles were observed to be in close proximity to each other (91±13 vesicles counted, depicted in Fig. S6B) and with varying diameter (ranging from 1 to 7 µm), but upon overnight tumbling at 25 ºC, they dispersed homogenously in solution (Fig. 3C,D).

Encapsulation of HPTS

10 µL of a 10 mM solution of pure phospholipid **3** in MeOH/$CHCl_3$ (1:1) was added to a glass vial, placed under a steady flow of $N_2$, and dried for 10 min to prepare a lipid film. Then, 100 µL of 0.1 mM HPTS aqueous solution was added to the lipid film and briefly vortexed. The solution was tumbled at room temperature for 30 min. Afterward, the resulting cloudy solution was diluted with an additional 200 µL of $H_2O$ and transferred to a 100 kDa molecular weight cut-off (MWCO) centrifugal membrane filter and centrifuged for 3 min at 10,000 rcf (Eppendorf 5415C). The solution was similarly washed for additional 5× to remove any nonencapsulated dye. Then, 1 µL of the vesicle solution were placed on a clean glass slide, secured by a greased cover slip, and imaged on a spinning disc confocal microscope (488 nm laser) to observe encapsulation of HPTS. Consistent with the previously described membrane-staining rehydration experiments, the phospholipid **3** in the presence of HPTS spontaneously self-assembled into vesicular structures with 0.5-8 µm diameter (163±24 vesicles counted). These HPTS-encapsulating

vesicles were observed to be in close proximity to each other (Fig. S6A), but upon overnight tumbling at 25 ºC, they dispersed homogenously in solution (Fig. 3B).

Transmission Electron Microscopy (TEM) Studies

**General.** A deposition System Balzers Med010 was used to evaporate a homogeneous layer of carbon. The samples were collected over 400 mesh Cu grids. The grids were then negatively stained with a solution of 1% (w/w) uranyl acetate. Micrographs were recorded on a FEI Tecnai™ Sphera microscope operating at 200 kV and equipped with a $LaB_6$ electron gun, using the standard cryotransfer holders developed by Gatan, Inc. For image processing, micrographs were digitized in a Zess SCAI scanner with different sampling windows.

**TEM measurements.** Copper grids (formvar/carbon-coated, 400 mesh copper) were prepared by glow discharging the surface at 20 mA for 1.5 min. Once the surface for vesicle adhesion is ready, 3.5 µL of a 5 mM solution of phospholipid **3** in $H_2O$ (previously hydrated at 37 ºC for 1 h) was deposited on the grid surface. This solution was allowed to sit for 10 seconds before being washed away with 10 drops of glass distilled $H_2O$ and subsequent staining with 3 drops of 1% w/w uranyl acetate. The stain was allowed to sit for 10 seconds before wicking away with filter paper. All grid treatments and simple depositions were on the dark/shiny/glossy formvar-coated face of the grid (this side face up during glow discharge). Samples were then imaged via TEM, revealing the presence of several populations of spherical compartments (50-950 nm in diameter), consistent with the vesicle architecture.

Phospholipid **3** formation from lysophospholipid **1** and commercially available palmitoyl-CoA

In a typical *de novo* phospholipid synthesis reaction, to a 5 µL of standard palmitoyl-CoA (10 mM stock in sterile $H_2O$), 5 µL of lysophospholipid **2** solution (10 mM stock in sterile $H_2O$) was successively added along with 40 µL of 100 mM $NaH_2PO_4/Na_2HPO_4$ buffer, pH 7.4 containing 10 mM TCEP, and the solution was mixed by gentle tapping. The reaction mixture was tumbled at 37 °C. HPLC-MS analysis was done after 4 h to analyze the formation of the corresponding product. Using UV traces at 205 nm, we observed that 820 µM phospholipid **3** was being made in the reaction. The product peak was comparable to the chemically synthesized phospholipid **3**.

Chemoenzymatic one-pot phospholipid formation mediated by cgFAS I

*In situ vesicle formation (without additives)*. For this reaction, lysophospholipid **2** was added to a scaled-up version of the cgFAS I-mediated palmitoyl-CoA (**1**) formation. In a typical *de novo* chemoenzymatic phospholipid synthesis reaction, to a 2 µL of lysophospholipid **2** solution (10 mM stock in sterile $H_2O$) was successively added 33 µL of 100 mM $NaH_2PO_4/Na_2HPO_4$ buffer, pH 7.4 containing 10 mM TCEP, along with 5 µL of 10 mM acetyl-CoA (in sterile $H_2O$), 5 µL of 100 mM NADPH (in sterile $H_2O$), and 5 µL of cgFAS (1 µM in elution buffer). Then, 5 µL of 70 mM malonyl-CoA (in sterile $H_2O$) was added, and the solution was mixed by gentle tapping. The reaction mixture was tumbled at 37 °C. Small aliquots (~2 µL) were taken out at various time points and placed on a glass slide for microscopic observations. Initially the vesicles were hard to visualize because of their small size, having sub-micrometer diameters. However, after tumbling overnight at 37 °C, some bigger vesicles were more evident, at 0.5-2 µm in diameter (133±12 vesicles counted, as indicated in Fig. 4C).

*In situ vesicle formation (with additives).* For the *de novo* phospholipid reaction with additives, the reaction was set up exactly as above, with the addition of 2.5 µL of guanidine hydrochloride (GuHCl) (10

mM in sterile $H_2O$), 2.5 µL of decanol (10 mM in sterile $H_2O$) and 2.5 µL of cholesterol (10 mM in EtOH) to the one-pot reaction. Cholesterol was added initially to the vial and $N_2$ gas was passed until all of the EtOH evaporated. Then, 2.5 µL of lysolipid **2** solution (10 mM in sterile $H_2O$) was added along with 25 µL of 100 mM $NaH_2PO_4$/$Na_2HPO_4$ buffer, pH 7.4 containing 10 mM TCEP, along with 5 µL of 10 mM acetyl-CoA (in sterile $H_2O$), 5 µL of 100 mM NADPH (in sterile $H_2O$), and 5 µL of cgFAS (1 µM in elution buffer). Then, 5 µL of 70 mM malonyl-CoA (in sterile $H_2O$) were added, and the solution was mixed by gentle tapping. The reaction mixture was tumbled at 37 °C. Small aliquots (~2 µL) were taken out at various time points and placed on a glass slide for microscopic observations. After 30 min, vesicles with 2.5-3 µm in diameter were evident in the reaction (Fig. 4D).

HPLC quantification of chemoenzymatic phospholipid **3** formation

Chromatographic separation was performed with a *Phase A/Phase B* gradients. A multistep gradient at a flow rate of 0.25 ml/min was used with the starting condition of *Phase B* at 5%, a linear increase to 50% until 7 min, then to 70% until 10 min and finally to 95% until 14 min runtime. For each HPLC-MS run, a small aliquot (10 µL) of the reaction was taken and then centrifuged to remove protein debris before loading onto the column. For the quantification, the UV traces at 205 nm were used (Fig. 4A,B), determining that the amount of phospholipid **3** made was 367 µM. The product peak was comparable to the chemically synthesized phospholipid **3**.

### 3. Supplementary Schemes

**A)**

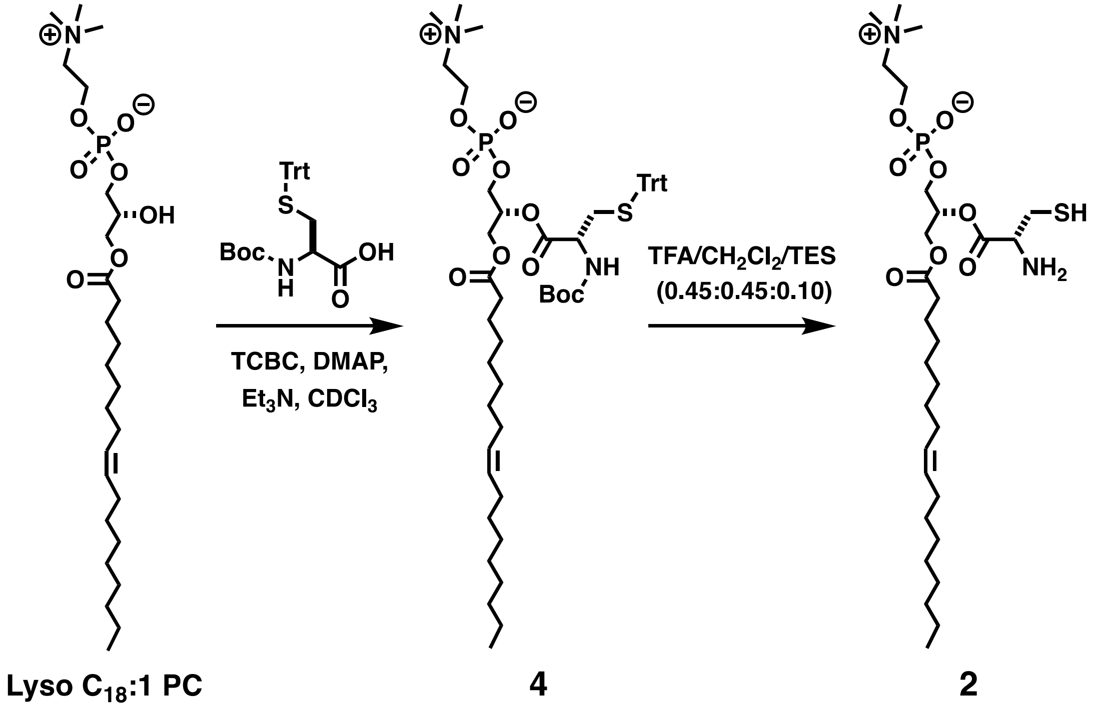


**B)**

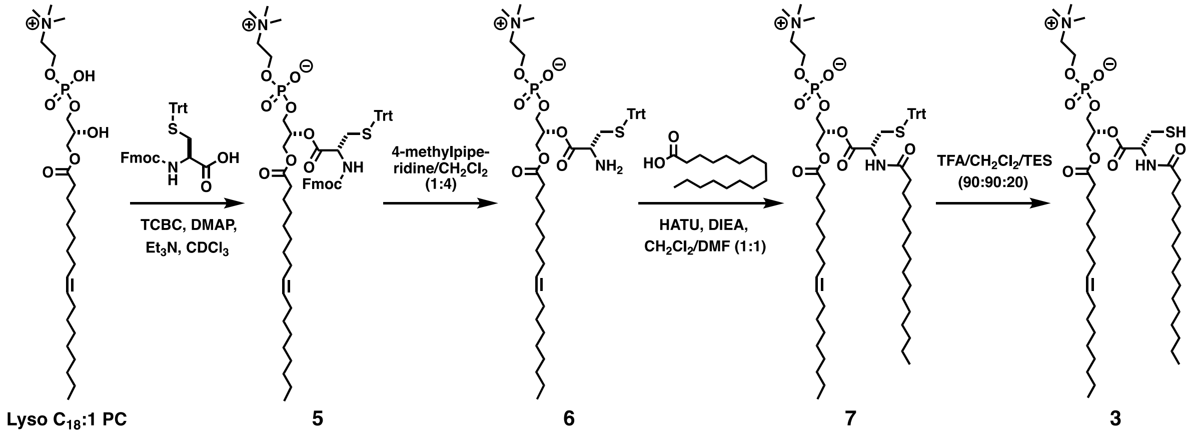


**Scheme S1.** Synthesis of lysophospholipid **2** (**A**) and phospholipid **3** (**B**).

### 4. Supplementary Figures

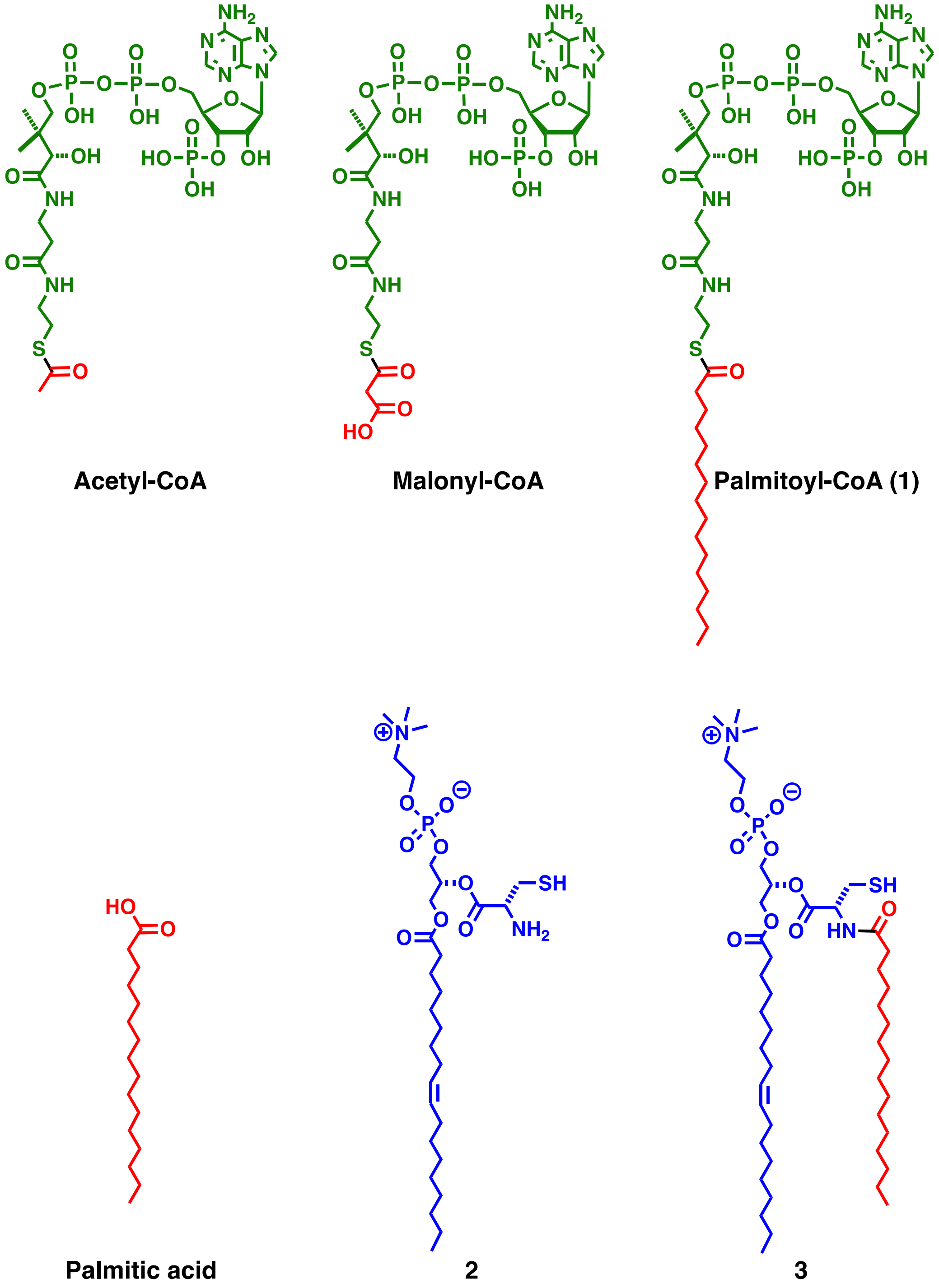


**Figure S1.** Chemical structures of all the lipids used in this study.

**A)**

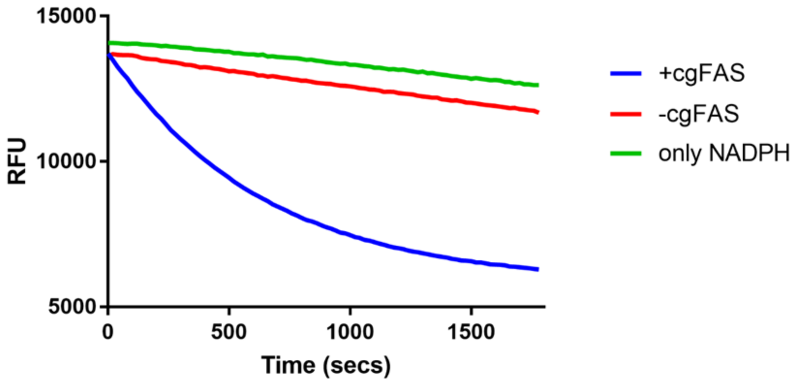


**B)**

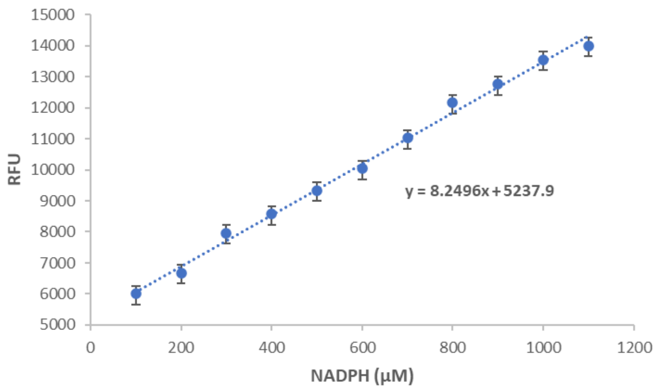


**C)**

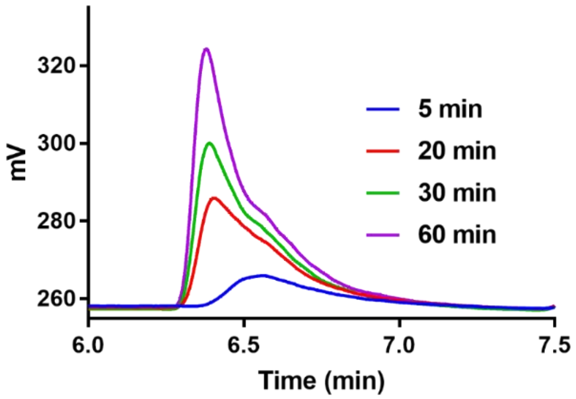


**Figure S2.** cgFAS-mediated palmitoyl-CoA (**1**) formation. **A)** Representative NADPH consumption assay. Fluorescence emission of NADPH is measured as the cgFAS-mediated palmitoyl-CoA formation consumes NADPH to convert it to $NADP^+$. **B)** Calibration curve for NADPH. **C)** HPLC-ELSD traces of palmitoyl-CoA formation over 1 h.

A)

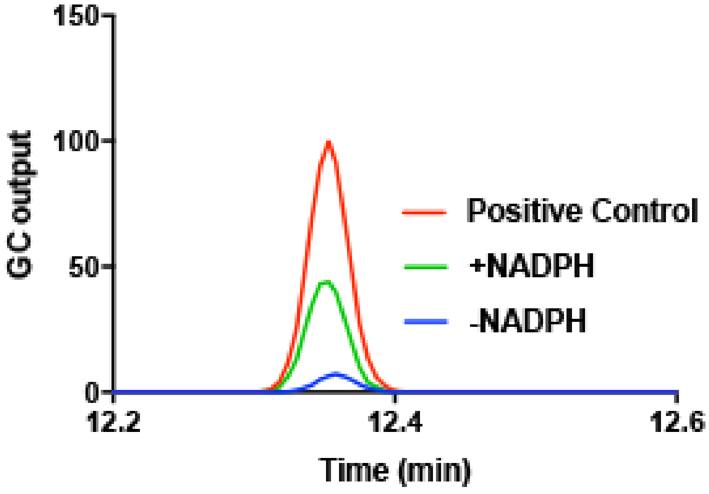


B)

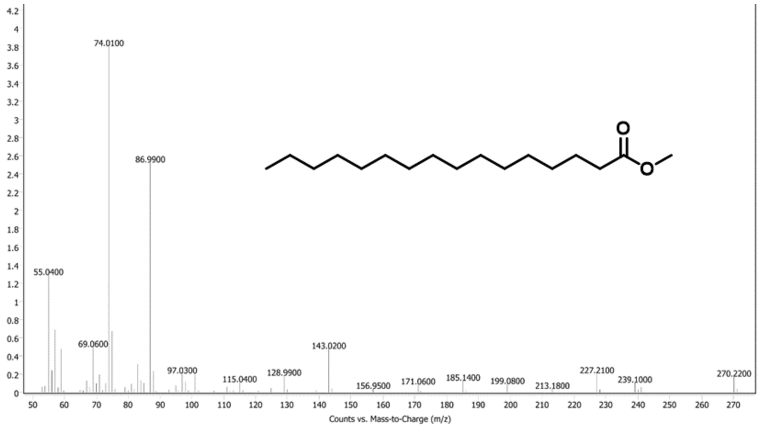


**Figure S3.** FAME analysis of cgFAS-mediated palmitoyl-CoA (**1**) formation. **A)** Representative FAME spectral analysis. The product is compared to the methyl ester of commercially available palmitoyl-CoA. **B)** Electronic impact (EI)-MS of the peak at 12.38 min, as methyl palmitate.

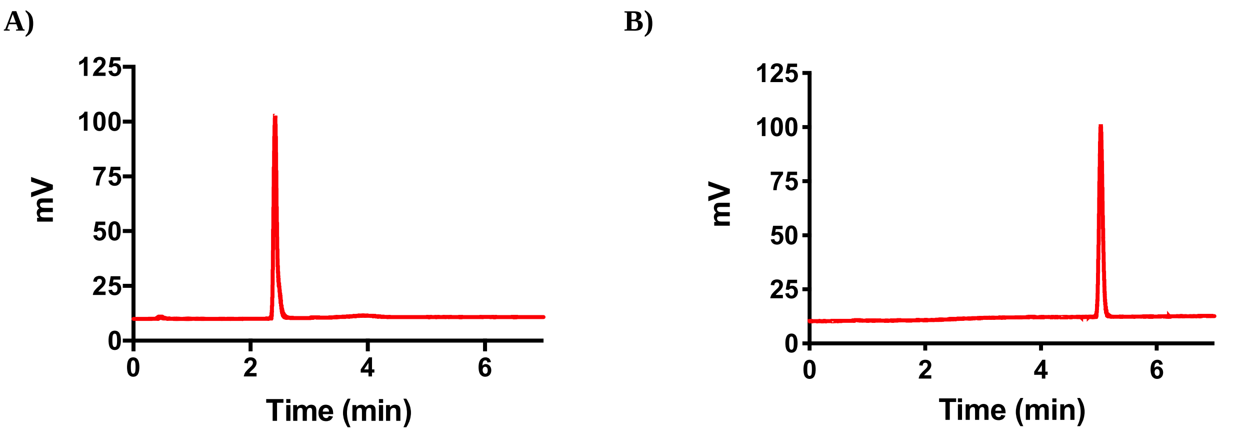


**Figure S4.** HPLC/ELSD spectra corresponding to lysophospholipid **2** (**A**) and phospholipid **3** (**B**). Retention times ($t_R$) were verified by mass spectrometry.

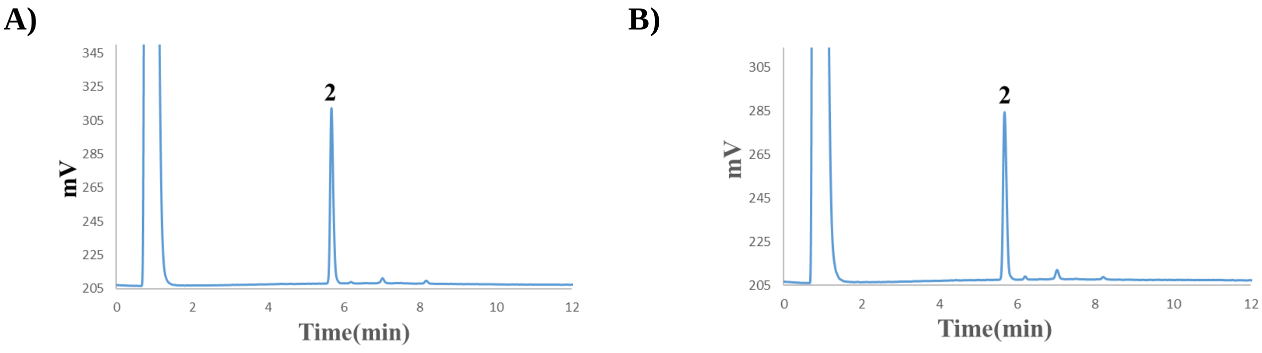


**Figure S5.** HPLC-ELSD traces corresponding to the reaction of FAS precursors (acetyl- and malonyl coA) with lysophospholipid **2** under standard NCL conditions (*negative controls*). Formation of ligation product was not observed for both cases, which highlights the importance of the hydrophobic interactions in the chemoselective reaction. **A)** NCL between acetyl-CoA and **2**. **B)** NCL between malonyl-CoA and **2**. Retention times ($t_R$) were verified by mass spectrometry.

A) B)

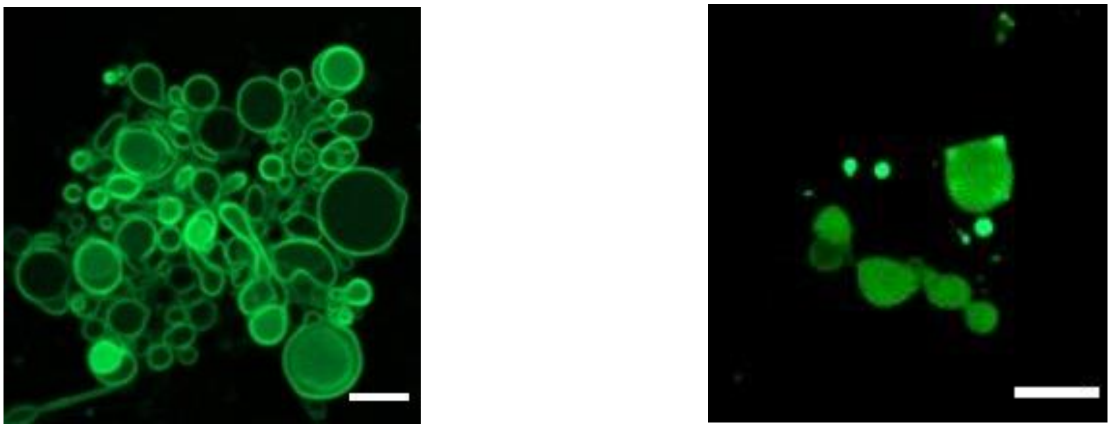

**Figure S6**. Spinning-disk confocal fluorescence microscope images demonstrating the spontaneous self-assembly of phospholipid **3** into membranous vesicles. **A)** Fluorescence microscopy image of vesicles of **3**. Membranes were stained with 0.1 mol % BODIPY-FL DHPE. **B)** Encapsulation of HPTS in membrane vesicles of **3**. Scale bars denote 5 μm.

A) B)

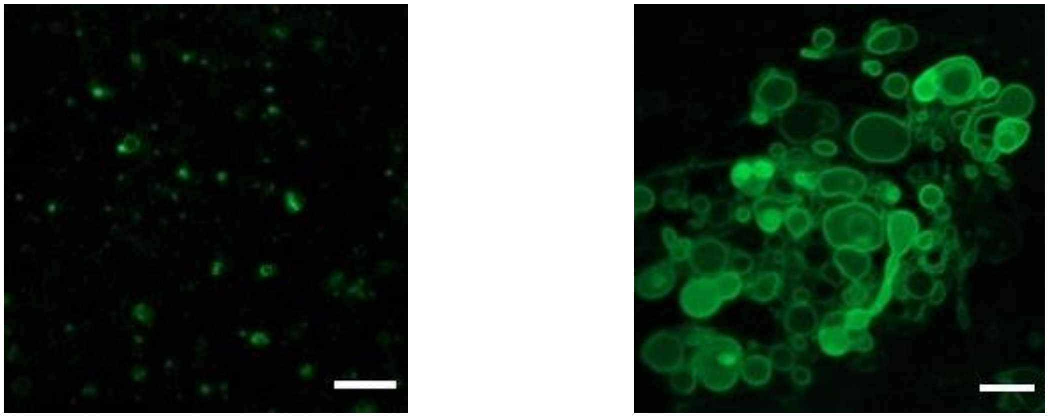

**Figure S7**. Spinning-disk confocal fluorescence microscope images demonstrating the chemoenzymatic *in situ* generation of phospholipid **3** and spontaneous formation of membranous vesicles. Fluorescence microscopy image of *in situ* chemoenzymatic reaction without **(A)** and with **(B)** 1:1:1 ratio of additives GuHCl, decanol and cholesterol. Membranes were stained with 0.1 mol % BODIPY-FL DHPE. Scale bars denote 5 μm.

### 5. NMR Spectra

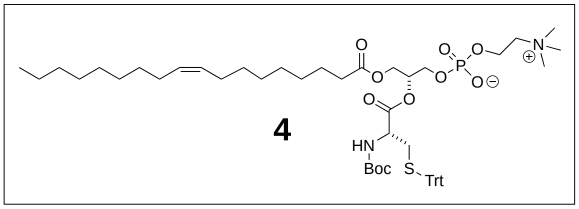


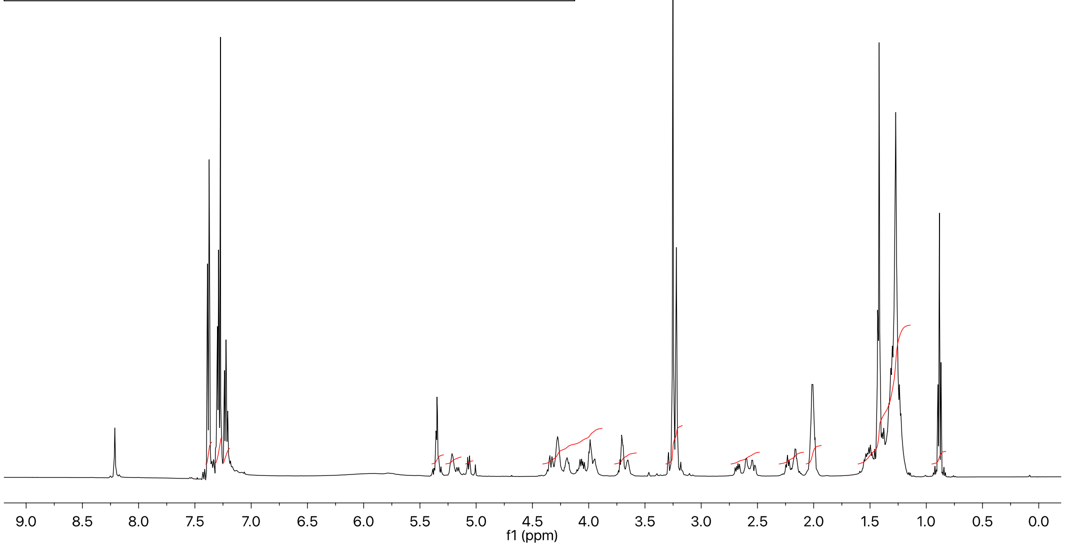


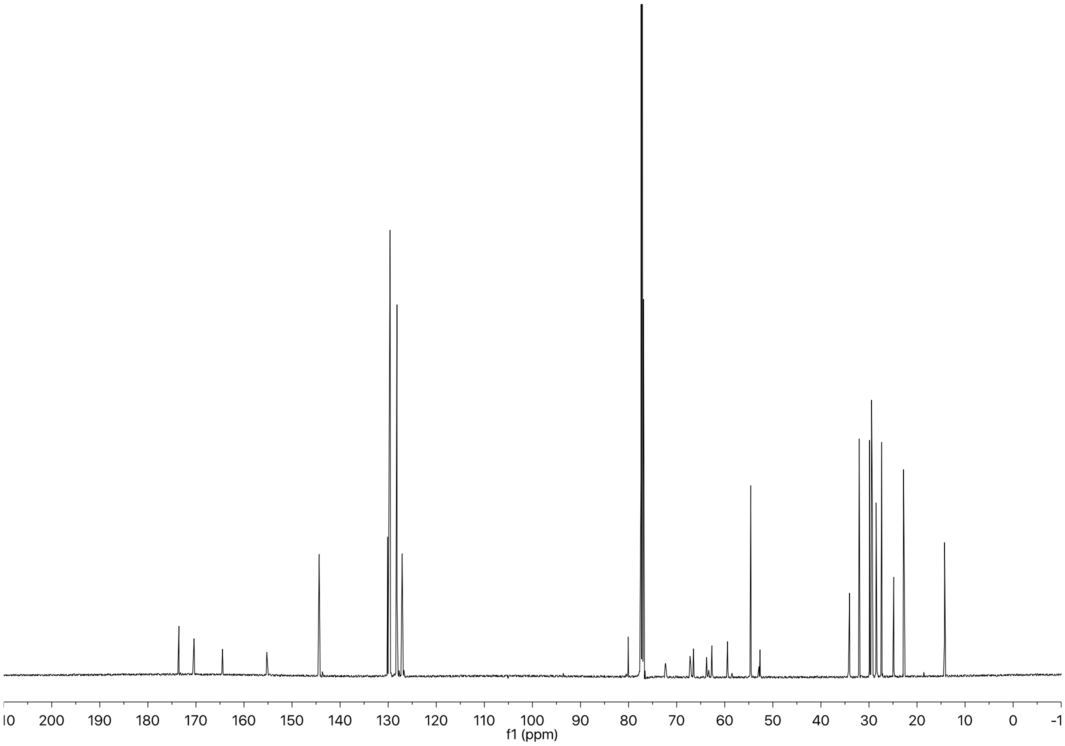

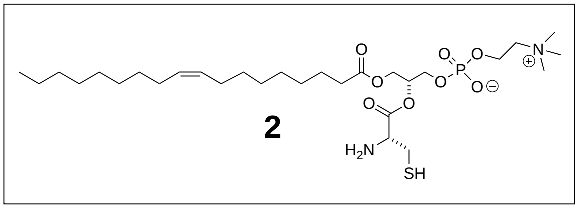

**2**

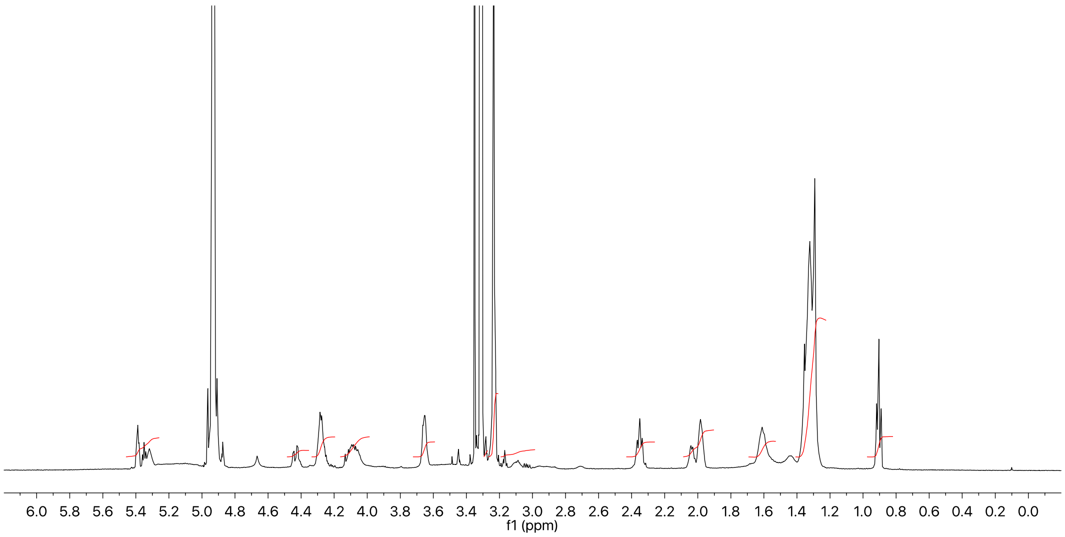
6.0 5.8 5.6 5.4 5.2 5.0 4.8 4.6 4.4 4.2 4.0 3.8 3.6 3.4 3.2 3.0 2.8 2.6 2.4 2.2 2.0 1.8 1.6 1.4 1.2 1.0 0.8 0.6 0.4 0.2 0.0
f1 (ppm)

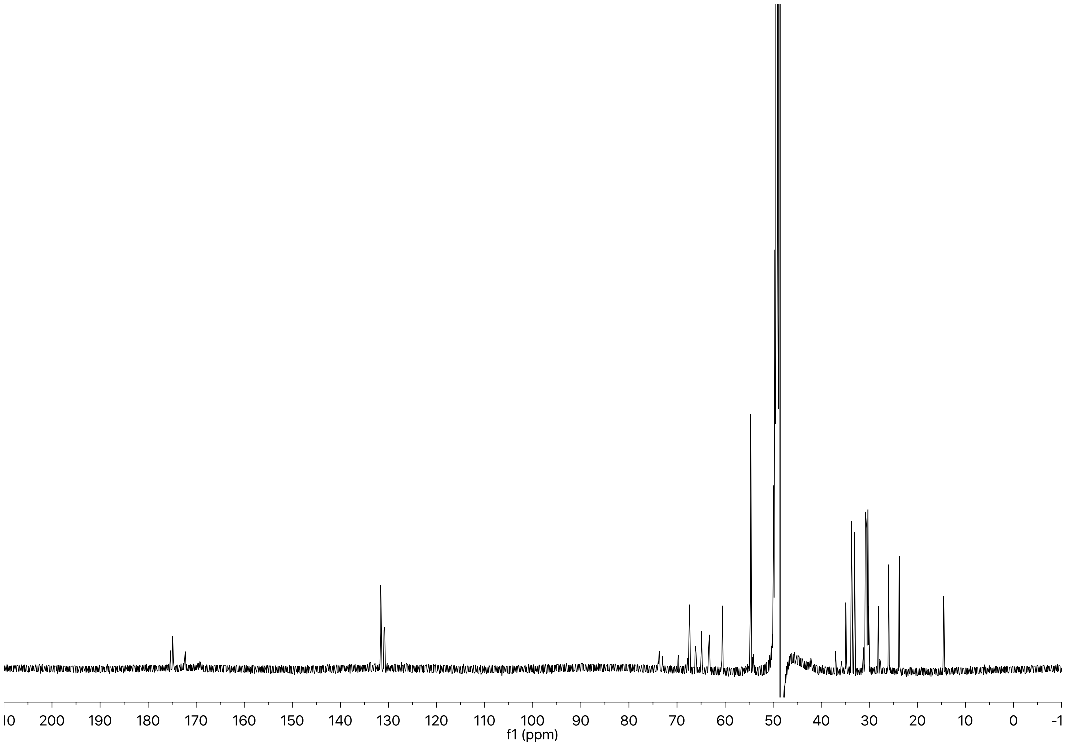
200 190 180 170 160 150 140 130 120 110 100 90 80 70 60 50 40 30 20 10 0
f1 (ppm)

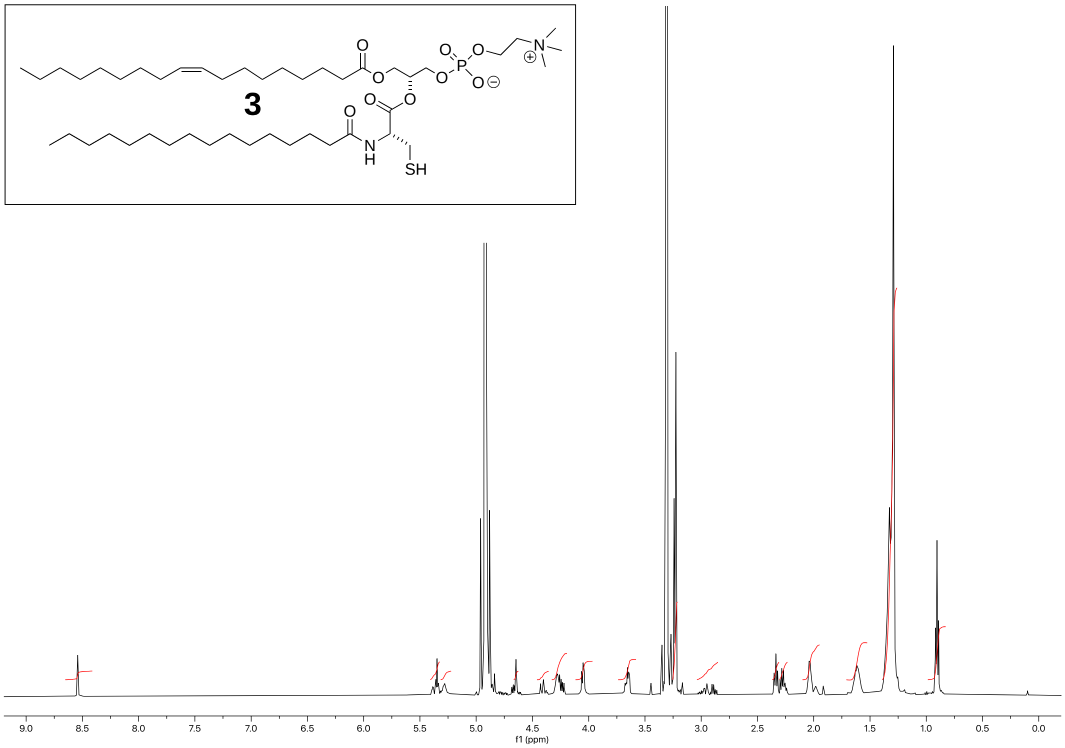
3
SH
9.0 8.5 8.0 7.5 7.0 6.5 6.0 5.5 5.0 4.5 4.0 3.5 3.0 2.5 2.0 1.5 1.0 0.5 0.0
f1 (ppm)

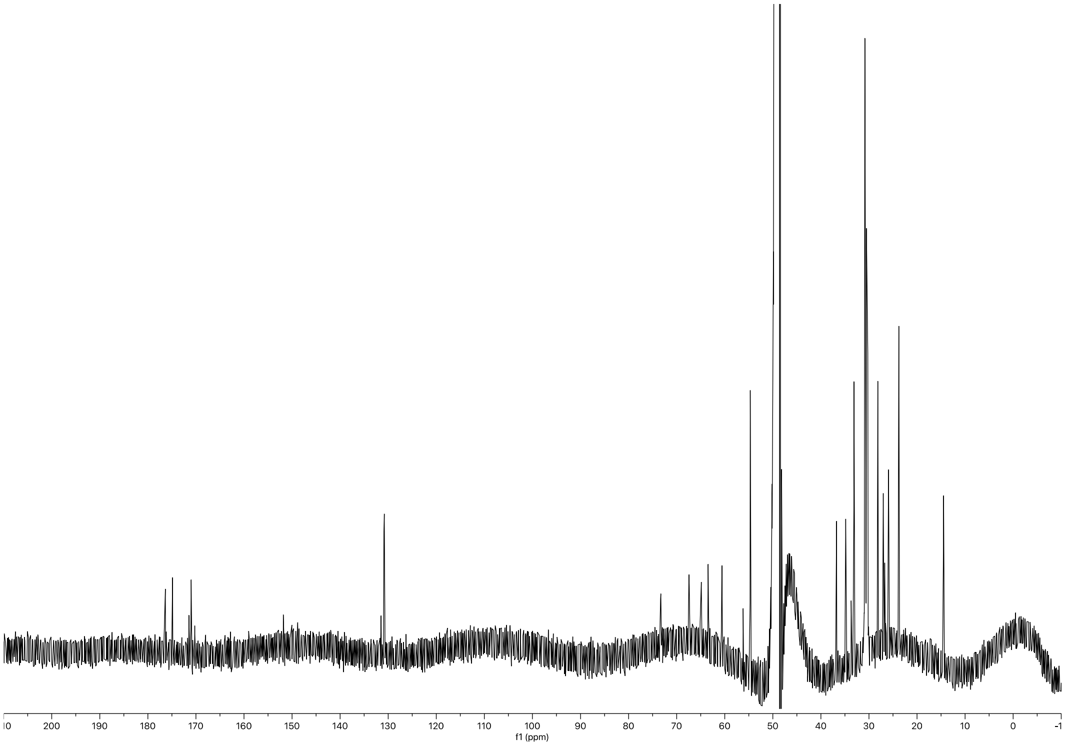
200 190 180 170 160 150 140 130 120 110 100 90 80 70 60 50 40 30 20 10 0
f1 (ppm)
